# Dissecting the dilemma between *in vitro* and *in vivo* drug screening: treating HepG2 cells with Artesunate as a model

**DOI:** 10.1101/499723

**Authors:** Johnny Kuan-Un Wong, Sophie Ling Shi, Haitao Wang, Fuqiang Xing, Yingyao Quan, Ming Zhao, Lei Zhang, Kristy Hio-Meng Wong, Ada Hang-Heng Wong, Chuxia Deng

## Abstract

The dilemma between *in vitro* and *in vivo* drug screening results persisted to hinder preclinical anti-cancer drug development. In this study, drug screening was initially performed on monolayer cultures of HepG2 cells, whereas *in vivo* drug testing was performed on the subcutaneous xenograft mouse model of HepG2 cells in athymic nude mice. Results showed that Artesunate inhibited HepG2 cell growth *in vitro*, but was ineffective *in vivo*. Hence, we investigated the difference between *in vitro* and *in vivo* settings using this cell-drug combination as a model. Immuno-staining of hepatocellular carcinoma (HCC) biomarkers showed that α-fetoprotein (AFP) was unaffected by Artesunate treatment *in vitro* or *in vivo*, suggesting that AFP was neither a biomarker nor response indicator. Alternatively, high albumin was detected in both monolayer and organoid cultures; contrariwise, the xenograft tumors prevailed low albumin, in consistency to human HCC with poor prognosis. Artesunate treatment reduced intracellular albumin expression *in vitro*; Artesunate did not alter tissue and serum albumin in xenograft mice, in coincidence with its irresponsiveness *in vivo*. However, Artesunate binding to albumin was undetectable. Instead, we observed a transient stimulation of Erk1/2 phosphorylation followed by DAPK1 dephosphorylation and apoptosis. Combined treatment of Artesunate with U0126 revoked Erk1/2-DAPK1 phosphorylation and exerted modest proliferative advantage at early time points, but eventually did not rescue HepG2 cells from death. U0126 induced multiploid formation independent of Artesunate, resulting in cell cycle arrest within 24 h post-treatment.

**Significance:** Systematic investigation of HCC biomarkers AFP and albumin among the *in vitro* models of monolayer and organoid cultures, and the *in vivo* models of subcutaneous and liver implantation xenograft mice was conducted.

## Introduction

The xenograft mouse model is the most prevalent preclinical model for anti-cancer drug development by far [1]. However, this method is costly, time-consuming and low-throughput. In contrast, *in vitro* drug screening provides a robust and cost-effective high-throughput alternative for lead compound identification and optimization. However, the unpredictable outcome *in vivo* results in substantial waste of resources. Many studies on closing the gap between *in vitro* and *in vivo* drug screening have been conducted. These include the development of organoid culture systems to mimick the three-dimensional structure of tumors [2], penetration of pseudo-blood vessels to mimick angiogenesis [3], construction of pseudo-liver and kidney modules to partially mimick the pharmacokinetics *in vivo* [4]. Nonetheless, the reproducibility between *in vitro* and *in vivo* drug screening results remained low. Since we are engaged in developing *in vitro* drug screening methods for precision cancer therapy [5], we are interested to delineate the dilemma between *in vitro* and *in vivo* drug response.

HepG2 is an immortalized human hepatocellular carcinoma (HCC) cell line. HCC is the most common form of liver cancer, which ranked third on the global cancer mortality list. Physical treatment of liver cancer includes hepatectomy, embolization and external beam radiation therapy (EBRT), whereas chemical treatment includes chemotherapy, targeted therapy and immunotherapy [6]]. Patients undergoing cancer therapy not only suffer from the disease, but also suffer from side effects, especially nausea, hair loss, *etc*, due to radiotherapy or chemotherapy. Hence, given that Traditional Chinese Medicine (TCM) is generally known to exert low cytotoxicity and mild side effects, we screened against some known to inhibit cancer in search for effective therapy.

Artesunate is a chemical derivative of Artemisinin. Artemisinin is a natural compound extracted from the herb *Artemisia annua* to combat malaria [7]. Later, Artemisinin and its derivatives were applied to other purposes, for example, Alzheimer’s disease [8], virus infections [9] and cancer [10–14]. Artemisinin and Artesunate underwent Phase I clinical trial to treat solid tumors caused by human papillomavirus. Artemisinin was reported to kill cancer cells by carbon-centered radicals generated by its endoperoxide bridge, the pathway of which was catalyzed by intracellular iron [12]. Artesunate, as a derivative of Artemisinin, was predicted to exert similar effects. However, this study showed that Artesunate did not induce apoptosis due to high intracellular iron in the liver cancer cell line HepG2. Artesunate did not trigger cell cycle arrest to inhibit HepG2 cell proliferation as reported in other cell lines too [11, 13]. In fact, screening of a total of eleven cancer cell lines prevailed that there was little correlation between intracellular iron and drug susceptibility. Instead, results showed that the Erk1/2-DAPK1 pathway was activated to trigger apoptosis *in vitro*. Erk1/2 phosphorylation is predominantly known to promote cell proliferation [15] and motility [16]. Death-associated protein kinase 1 (DAPK1) is a multi-domain protein kinase [17], phosphorylated by Erk1/2 at Ser735 to trigger apoptosis [18]. To test our hypothesis, we applied U0126, which is an inhibitor of the Erk1/2 upstream kinase MEK1 [19]. Nevertheless, although U0126 exerted cell proliferation advantage at early time points, it triggered multiploid formation independent of Artesunate, resulting in cell cycle arrest and eventually cell death.

## Results

### Artesunate inhibited HepG2 cell growth *in vitro* but was ineffective in subcutaneous xenograft mouse model

In this study, we screened eleven cancer cell lines belonging to six tissue origins against Artemisinin and its three derivatives, namely Artesunate, Dihydroartemisinin and Artemether. Results showed that Artesunate was the most potent Artemisinin analog *in vitro* (Figure 1A). Moreover, the hepatocellular carcinoma (HCC) cancer cell line HepG2 was susceptible to all four analogs (Figure 1B), whereas the cholangiocarcinoma (CC) cancer cell line QBC939 was susceptible to Artesunate and Dihydroartemisinin only (Figure 1C).

**Figure 1.**
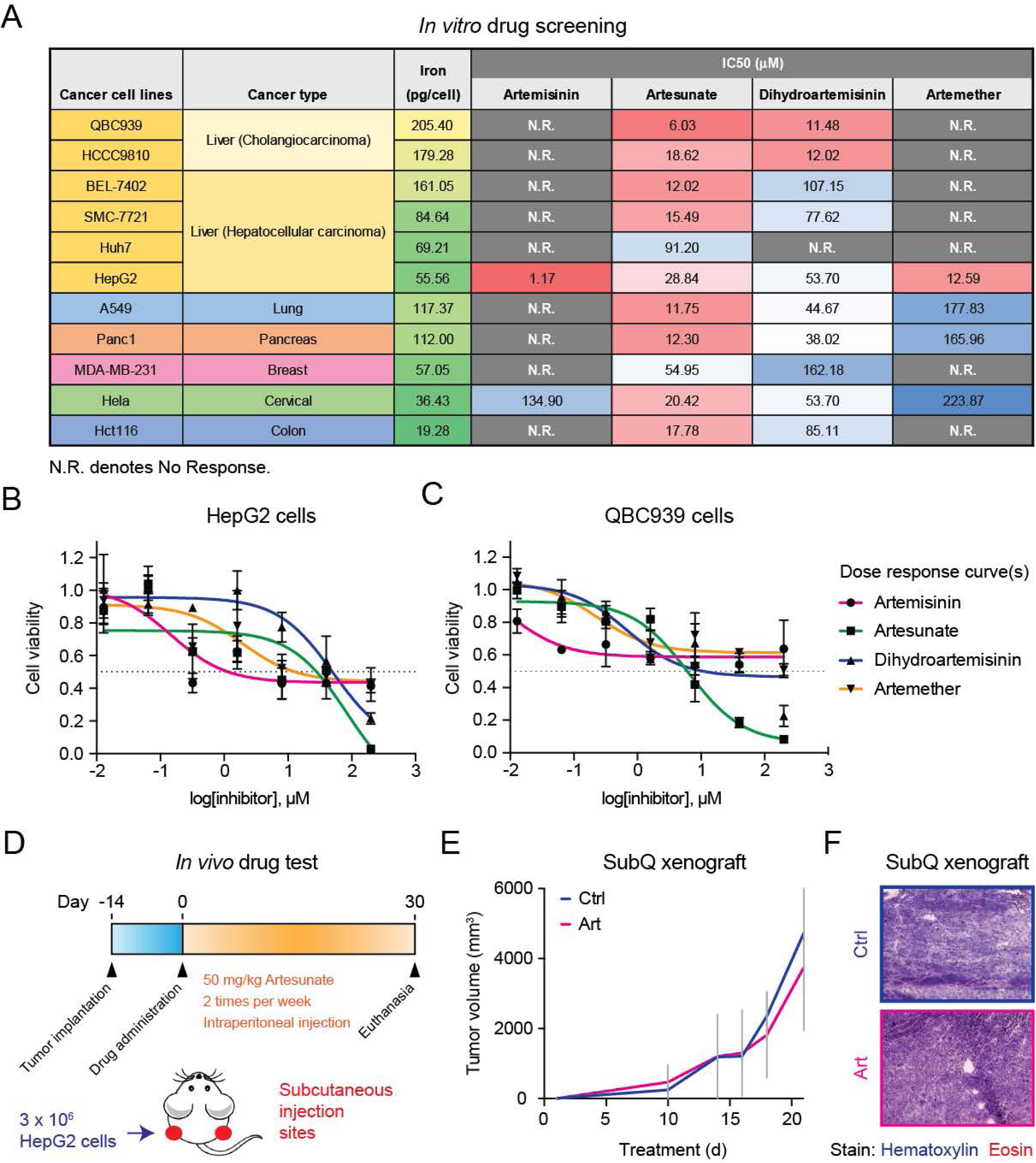
Artesunate was effective *in vitro* but ineffective *in vivo*. (A) Drug screening and intracellular iron measurement of cancer cell lines. A sum of eleven cancer cell lines from six tissue origins were screened against four analogs of Artemisinin. Cell viability was assessed by Alamar Blue assay and intracellular iron was measured by ionization coupled plasma atomic emission spectroscopy. Quadriplicates were used for drug screening and single point measurement was used for intracellular iron measurement. (B) Drug response curves of HepG2 treated with four Artemisinin analogs. HepG2 cells were treated with gradient doses of Artemisinin and its derivatives from 200 μM to 12.8 nM for 3 d, and subject to Alamar Blue assay. The mean and standard deviation of normalized cell viability from quadruplicate experiments was plotted against the log of drug concentration. (C) Drug response curves of QBC939 treated with four Artemisinin analogs. HepG2 cells were treated with gradient doses of Artemisinin and its derivatives from 200 μM to 12.8 nM for 3 d, and subject to Alamar Blue assay. The mean and standard deviation of normalized cell viability from quadruplicate experiments was plotted against the log of drug concentration. (D) Subcutaneous xenograft mouse model of HepG2 cells. HepG2 cells was subcutaneously implanted in athymic nude mice and administered with 50 mg/kg Artesunate two times per week by intraperitoneal injection for 30 d; parallel treatment with solvent alone was performed in control mice. (E) Drug efficacy evaluation by tumor volume. Tumor size of the subcutaneous xenograft mice was measured at indicated time intervals and plotted on graph. The mean and standard deviation of tumor volumes of all replicate mice in each treatment group were plotted against treatment time. (F) H&E staining of xenograft tumors. Tumors were resected, fixed and stained by hematoxylin and eosin after euthanasia, followed by brightfield imaging under 4x objective.

Next, we tested the treatment efficiency of Artesunate in a subcutaneous xenograft mouse model. HepG2 cells were subcutaneously implanted into the left and right flank of athymic nude mice (Figure 1D). When tumors grew to 2 mm in any direction, 50 mg/kg Artesunate was intraperitoneally injected to treat the mice. However, tumors continued to grow after Artesunate treatment *in vivo* (Figure 1E). Hematoxylin and eosin (H&E) staining of the xenograft tumors showed that there was no significant difference between control and Artesunate-treated mice (Figure 1F), indicating that Artesunate was ineffective in inhibiting tumor growth *in vivo*.

It is prominent in preclinical anti-cancer drug development that potent drugs *in vitro* could be ineffective *in vivo*. Therefore, we analyzed the HCC biomarkers of α-fetoprotein (AFP) and albumin in monolayer and organoid HepG2 cultures *in vitro*, in comparison to HepG2-engrafted tumors in subcutaneous and liver implantation xenograft models, respectively. No Artesunate treatment was conducted in the liver implantation xenograft model. In parallel experiments to the xenograft models, untreated nude mice that was not engrafted nor treated, was used to compare with HepG2-engrafted mice under mock treatment.

### AFP was neither a biomarker nor response indicator in our comparison model

Immunofluorescence (IF) staining of AFP showed that HepG2 cells expressed AFP on monolayer cultures *in vitro*, with similar expression levels between control and Artesunate-treated cells (Figure 2A). However, AFP expression was compromised in organoid culture *in vitro*, regardless of treatment (Figure 2B), suggesting that AFP expression could be directly regulated by tissue architecture. Similar to the organoids, immunohistochemical (IHC) staining of AFP was negative on the epithelial regions of the subcutaneous implanted tumors in both control and Artesunate-treated mice (Figure 2C). Low AFP expression was also observed in engrafted tumors of the liver implantation xenograft model (Figure 2D). In summary, AFP was only highly expressed in monolayer HepG2 cultures, but lowly expressed in organoid cultures and the xenograft tumors. Additionally, AFP expression was unaffected by Artesunate treatment *in vitro* and *in vivo*.

**Figure 2.**
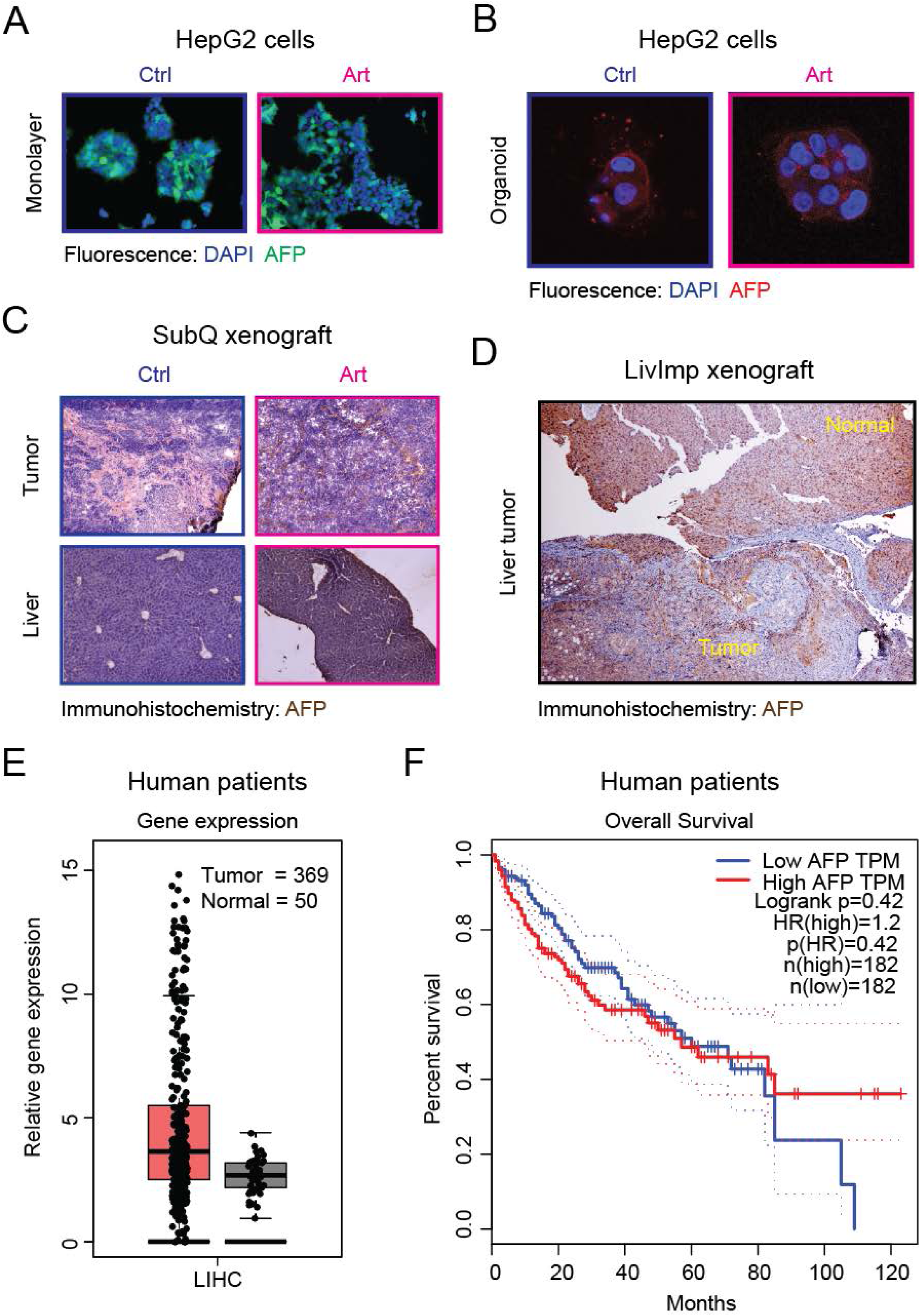
The HCC biomarker AFP was analyzed. (A) IF staining of AFP in HepG2 monolayer cultures after Artesunate treatment. HepG2 cells were treated in the absence and presence of 30 μM Artesunate for 12-18 h, followed by formalin fixation and AFP antibody staining. Fluorescence images were taken under 20x objective. (B) IF staining of AFP in HepG2 organoid cultures after Artesunate treatment. HepG2 cells were treated in the absence and presence of 30 μM Artesunate for 12-18 h, followed by formalin fixation and AFP antibody staining. Fluorescence images were taken under 60x objective. (C) IHC staining of AFP in subcutaneous xenograft mice. Tumors and livers of control and Artesunate-treated subcutaneous xenograft mice were fixed and stained with AFP antibody. Brightfield images were taken using 4x objective. (D) IHC staining of AFP in liver implantation xenograft mice. Tumors and adjacent liver tissue of untreated liver implantation xenograft mice were fixed and stained with AFP antibody. Brightfield images were taken using 4x objective. (E) Gene expression profile analysis of AFP in clinical cohorts by Gene Expression Profiling Interactive Analysis (GEPIA; http://gepia.cancer-pku.cn/). AFP expression between cancerous and normal tissue in HCC patients was shown. (F) Overall survival curves of liver cancer patients with different levels of AFP expression by GEPIA.

Next, we looked up the database for AFP expression in human HCC patients and its relationship to disease prognosis. Analysis of AFP mRNA expression in human HCC patients showed insignificant difference between cancerous and paired normal tissue, with substantial variance in cancerous tissue (Figure 2E). In concert to human mRNA expression, engrafted tumors and liver hepatocytes of the subcutaneous xenograft model prevailed low AFP expression (Figure 2C). Nevertheless, liver hepatocytes depicted high AFP expression as compared to low AFP in adjacent tumor in our liver implantation xenograft model (Figure 2D). This observation was consistent with the high variance of AFP mRNA in cancerous tissue (Figure 2E), and reinforced the notion that AFP was elevated in some but not all HCC patients [20]. Lastly, AFP levels was uncorrelated to patients’ survival (Figure 2F).

Taken together, AFP was merely highly expressed in monolayer HepG2 cells and in liver hepatocytes in the liver implantation model. This suggested that tumors might not produce AFP. Instead, liver cancer might trigger hepatocytes to produce AFP through proximal interaction, because the liver implantation xenograft mice but not the subcutaneous xenograft mice exhibited elevated AFP in liver hepatocytes (Figure 2C and 2D). Consistently, untreated nude mice in both xenograft models depicted low intracellular AFP (Supplementary Figure 1A). Consequently, we deduced that AFP was neither a biomarker nor response indicator in our comparison model.

### Reduction in intracellular albumin was a response indicator *in vitro*

Albumin depicted positive IF staining in monolayer HepG2 cells, and diminished after Artesunate treatment (Figure 3A). HepG2 organoids demonstrated similar phenomenon to monolayer cultures (Figure 3B). In contrary to *in vitro* cultured cells, intracellular albumin was low in subcutaneous xenograft tumors regardless of treatment (Figure 3C). Consistently, low albumin was detected in xenograft tumors of the liver implantation model (Figure 3D). In contrast to the implanted tumors, albumin was highly expressed in liver hepatocytes in both the subcutaneous and liver implantation models (Figure 3C and 3D). In summary, albumin was highly expressed in both monolayer and organoid cultures *in vitro*, but lowly expressed in xenograft tumors *in vivo*. Moreover, Artesunate reduced albumin in *in vitro* cultures, but had no impact on tissue albumin in our subcutaneous xenograft model.

**Figure 3.**
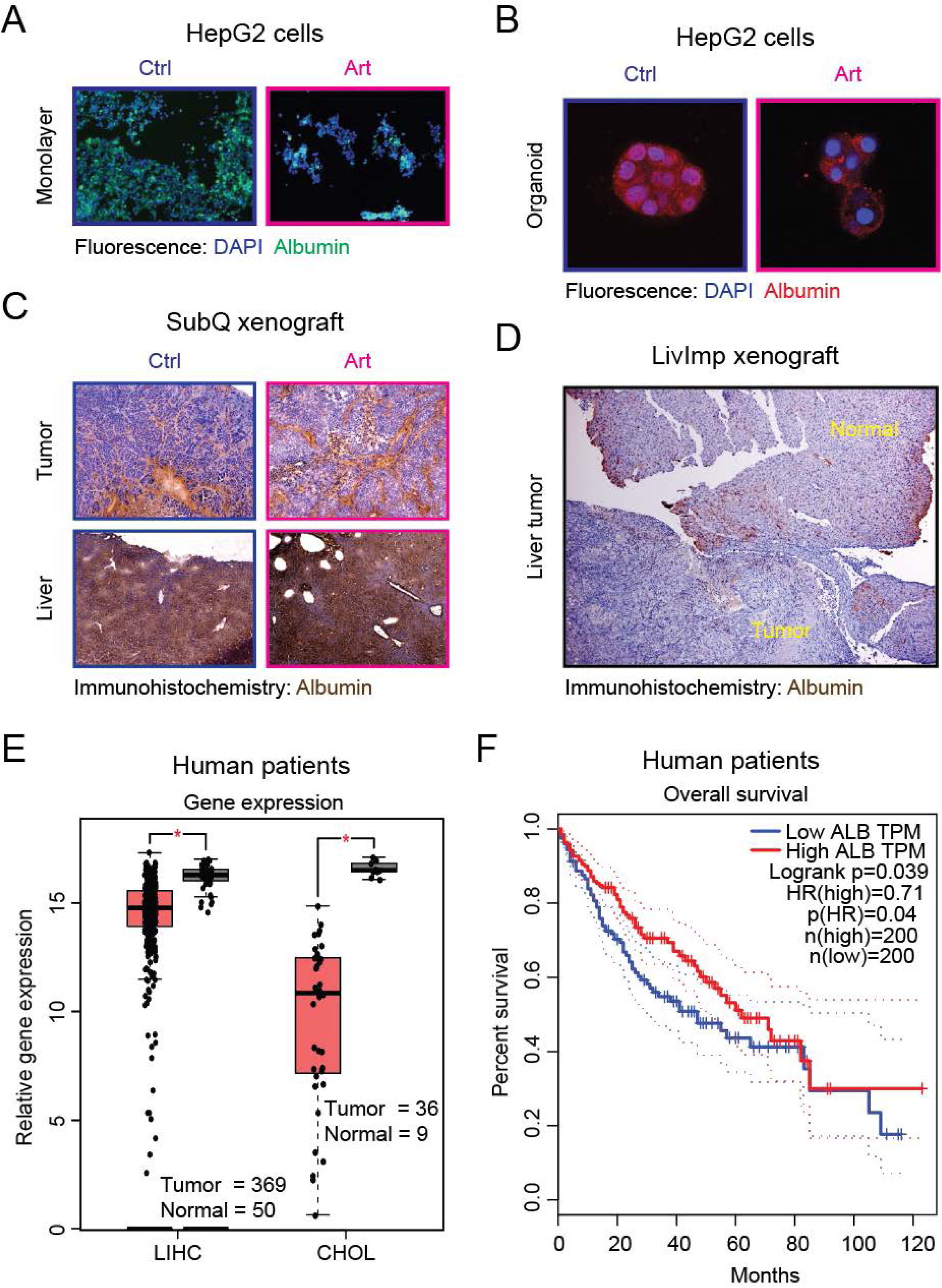
The plasma-abundant protein albumin was analyzed. (A) IF staining of albumin in HepG2 monolayer cultures after Artesunate treatment. HepG2 cells were treated in the absence and presence of 30 μM Artesunate for 12-18 h, followed by formalin fixation and albumin antibody staining. Fluorescence images were taken under 20x objective. (B) IF staining of albumin in HepG2 organoid cultures after Artesunate treatment. HepG2 cells were treated in the absence and presence of 30 μM Artesunate for 12-18 h, followed by formalin fixation and albumin antibody staining. Fluorescence images were taken under 60x objective. (C) IHC staining of albumin in subcutaneous xenograft mice. Tumors and livers of control and Artesunate-treated subcutaneous xenograft mice were fixed and stained with albumin antibody. Brightfield images were taken using 4x objective. (D) IHC staining of albumin in liver implantation xenograft mice. Tumors and adjacent liver tissue of untreated liver implantation xenograft mice were fixed and stained with albumin antibody. Brightfield images were taken using 4x objective. (E) Gene expression profile analysis of albumin (ALB) in clinical cohorts by Gene Expression Profiling Interactive Analysis (GEPIA; http://gepia.cancer-pku.cn/). ALB expression between cancerous and normal tissue in HCC and CC patients. (F) Overall survival curves of liver cancer patients with different levels of ALB expression by GEPIA.

In concert to the IHC staining results, lower albumin mRNA expression was observed in cancerous tissue as compared to paired normal tissue in both human HCC and CC patients (Figure 3E). Low albumin level was correlated to poor prognosis in humans surviving below 70 months after confirmed diagnosis of liver cancer (Figure 3F).

Next, we measured secreted albumin in our comparison model. Firstly, both the subcutaneous and liver implantation xenograft mice demonstrated no statistical difference in blood serum albumin as compared to untreated nude mice (Figure 4A). Secondly, the subcutaneous xenograft mice treated with Artesunate depicted no statistical difference in blood serum albumin as compared to control (Figure 4A). Thirdly, albumin was undetectable in monolayer HepG2 cells regardless of treatment (Figure 4B). Fourthly, HepG2 organoids secreted albumin at similar levels under both control and Artesunate-treated conditions (Figure 4B). This finding demonstrated that Artesunate merely reduced intracellular albumin but not secreted albumin *in vitro*. The lack of response in secreted albumin after Artesunate treatment might be explained by the slow synthesis of approximately 8 μg h^-1^ plate^-1^ [21] and the long half-life of 3 weeks [22] of albumin, so that disruption of its neosynthesis could not be detected in culture medium within 12 h post-treatment. Moreover, predominant expression of albumin in many murine organs (Figure 4C) but not the engrafted tumors suggested that serum albumin could not serve as a response indicator *in vivo*.

**Figure 4.**
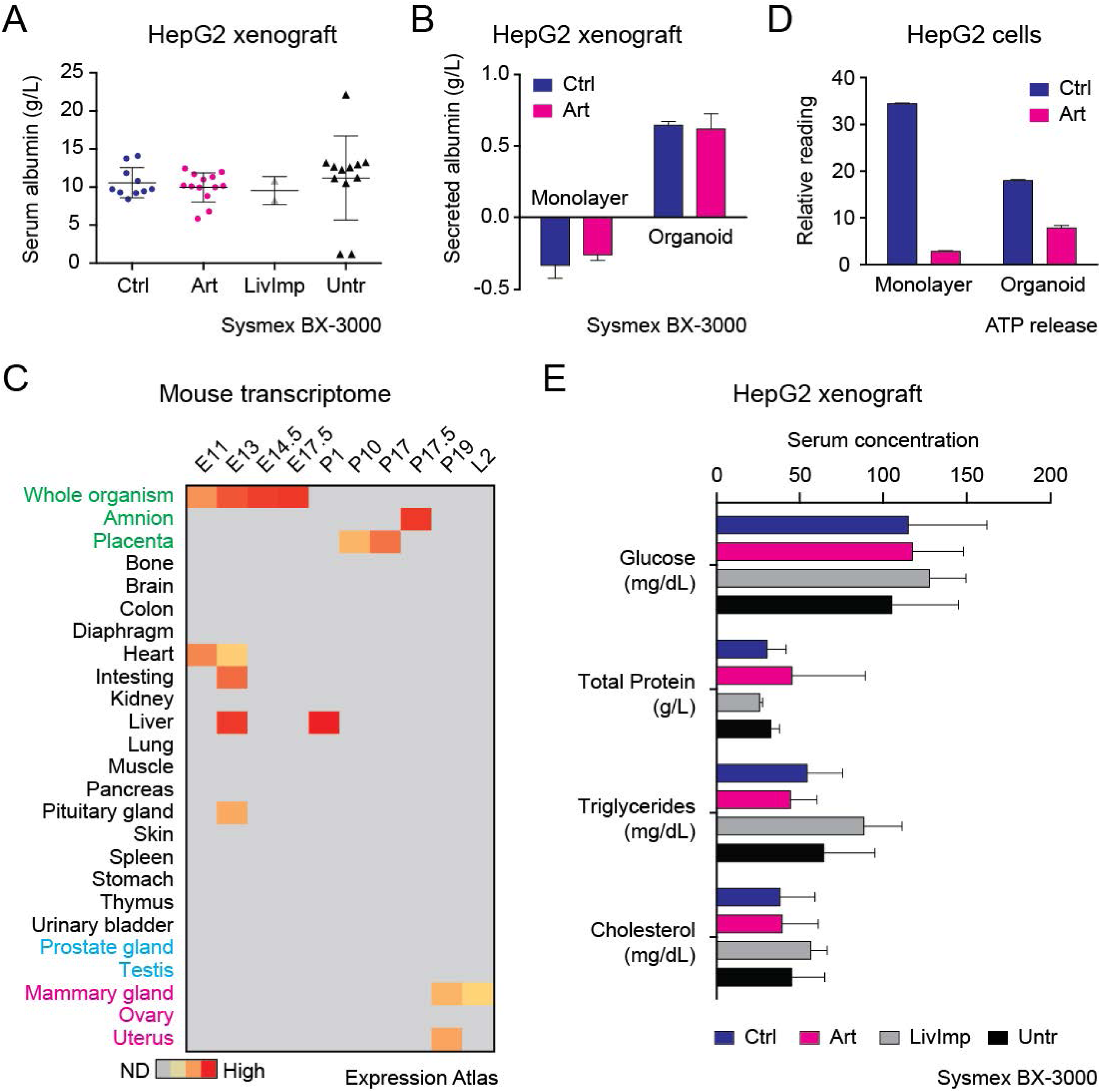
Secreted albumin from the *in vitro* and *in vivo* models was analyzed. (A) Blood biochemistry of HepG2 xenograft mice. Blood serum from control and Artesunate-treated subcutaneous xenograft mice, and liver implantation xenograft mice, was extracted and subject to biochemical analysis for albumin concentration; blood serum from untreated nude mice was measured in parallel. (B) Secreted albumin measurement in HepG2 monolayer and organoid cultures after Artesunate treatment. Serum-depleted culture medium was incubated with HepG2 cells on monolayer and in organoid cultures under control and Artesunate treatment for 12 h, and subject to biochemical analysis for albumin concentration (g/L). Albumin concentration detected from serum-depleted culture medium was used as blank. (C) Analysis of albumin mRNA expression by Expression Atlas (https://www.ebi.ac.uk/gxa/). Partial mRNA results from 49 Fantom5 Project depicting different developmental stages of *Mus musculus* was used. All values were logged by 10, followed by a red-yellow color coding of mRNA expression from high to low; grey color indicated NA in raw data. (D) Cell proliferation assay of HepG2 monolayer and organoid cultures after Artesunate treatment. 30 μM Artesunate was applied to HepG2 cells on monolayer or in organoid cultures for 2 d, and subject to ATP release assay following manufacturer’s protocol. (E) Blood biochemical analysis of HepG2 xenograft mice. Blood serum from control or Artesunate-treated HepG2 xenograft mice, and liver implantation xenograft mice, was extracted and subject to biochemical analysis for absolute concentrations of glucose (mg/dL), total protein (g/L), triglycerides (mg/dL) and cholesterol (mg/dL); blood serum from untreated nude mice was measured in parallel.

To verify that Artesunate treatment was effective in both *in vitro* models, cell proliferation of monolayer and organoid HepG2 cells was measured by ATP release assay. Results showed that proliferation of both HepG2 monolayer and organoid cultures treated with Artesunate slowed down as compared to corresponding controls (Figure 4D), suggesting that Artesunate was effective in both monolayer and organoid cultures. Notably, monolayer culture enabled faster cell proliferation than organoid culture (Figure 4D), hence dampening the proliferation retardation in Artesunate-treated organoids as compared to monolayer cells (Figure 4D). However, because intracellular albumin could not be conclusively quantified, we could not deduce where there was proportionate relationship between Artesunate efficacy and intracellular albumin reduction *in vitro*. Nevertheless, these data indicated that tissue architecture alone could not explain the inefficacy of Artesunate *in vivo*.

Lastly, we checked the blood biochemical parameters of xenograft mice in comparison to untreated nude mice, to investigate the impact of tumor engraftment and drug administration on physiology. Results showed that there was insignificant difference in glucose, total protein, triglycerides and cholesterol among the subcutaneous xenograft model, the liver implantation model and untreated mice (Figure 4E). Artesunate treatment did not significantly alter these parameters as compared to control in subcutaneous xenograft mice (Figure 4E). Body weight of the different experimental groups exhibited no significant difference too (data not shown). Hence, we concluded that the investigated animal blood biochemical parameters were insignificantly altered by HepG2 engraftment or Artesunate treatment.

Albumin frequently serves as a drug carrier *in vivo* [23]. Thus, we first tested if albumin binds to Artesunate to enhance intracellular uptake of Artesunate in *in vitro* cultures. Molecular docking of Artesunate on human albumin protein predicted three binding sites (Supplementary Figure 2A). However, no binding between Artesunate and bovine serum albumin was observed up to 4.25:1 molar ratio between Artesunate and albumin using isothermal calorimetry (ITC) (Supplementary Figure 2B). To verify that our comparisons were valid, sequence alignment of human, bovine and mouse albumin proteins was performed. Results showed that the predicted binding sites were generally conserved across species (Supplementary Figure 2C), suggesting that the ITC binding experiment was relevant to imply for mouse albumin in our xenograft model. Hence, we concluded that Artesunate did not bind to albumin.

Based on all prevailing evidence, albumin seemed to be consequence rather than cause of effective Artesunate treatment *in vitro*, suggesting that albumin acts as a response indicator of Artesunate treatment *in vitro*.

### Artesunate inhibited HepG2 cell proliferation and activated Erk1/2-DAPK1 pathway to promote apoptosis *in vitro*

After that, we sought to dissect the mechanism of Artesunate in inhibiting HepG2 cell proliferation. Initially, we tested known Artemisinin modes of action. Firstly, even though liver is rich in iron (Supplementary Figure 3A), not all liver cancer cell lines depicted high intracellular iron concentration (Figure 1A and Supplementary Figure 3B). Indeed, HepG2 cells contained the lowest intracellular iron among all liver cancer cell lines tested in this study (Figure 1A and Supplementary Figure 3B). Nevertheless, HepG2 cells still exhibited low Artesunate IC_50_ as compared to other cell lines (Figure 1A and Supplementary Figure 3B). Secondly, we did not see significant cell cycle arrest in HepG2 cells treated with Artesunate as compared to control at 12 h post-treatment (Supplementary Figure 3C), suggesting that Artesunate did not trigger cell cycle arrest to inhibit HepG2 cell proliferation. Thirdly, radical oxygen species (ROS) staining did not show significant increase in HepG2 cells treated with any of the four Artemisinin analogs at 12 h post-treatment (Supplementary Figure 3D), in consistence with the low intracellular iron in HepG2 cells (Figure 1A and Supplementary Figure 3B). Consistently, there was no significant pH changes in HepG2 cells after 12 h treatment with Artesunate as compared to control (Supplementary Figure 3E). Taken together, our results depicted that Artesunate inhibited HepG2 cell proliferation without compliance to known Artemisinin modes of action.

In order to decipher the mechanism of action of Artesunate on HepG2 cells, we tested the phosphorylation of three kinases, namely Erk1/2, p38 and AMPKα, after Artesunate addition to HepG2 cells by Western blotting. Western blot analysis showed that phosphorylated Erk1/2 decreased at 1 h post-treatment in Artesunate-treated HepG2 cells (Supplementary Figure 4A). On the other hand, increased phosphorylated p38 was observed at 10 min post-treatment in control and Artesunate-treated HepG2 cells respectively (Supplementary Figure 4A). No AMPKα phosphorylation was observed in control and Artesunate-treated HepG2 cells (Supplementary Figure 4A). Hence, we deduced that Erk signaling was involved in HepG2 cytotoxicity by Artesunate. Indeed, U0126 alleviated the cytotoxic effect of Artesunate treatment (Supplementary Figure 4B).

Next, we applied U0126 to HepG2 cells in the absence or presence of Artesunate. First, we observed elevated Erk1/2 phosphorylation in different patterns in the absence and presence of medium replacement during Artesunate addition (Figures 5A and 5B). In both scenarios, DAPK1 dephosphorylation occurred at 30 min post-treatment (Figures 5A and 5B). In contrast, DAPK1 dephosphorylation was unapparent in HepG2 cells treated with U0126 alone or in combination with Artesunate without medium replacement (Figure 5A), whereas DAPK1 dephosphorylation was observed in control, U0126 and double treatment at 30 min after medium replacement (Figure 5B), suggesting of slightly different Erk1/2-DAPK1 signaling in the absence and presence of medium replacement that was independent of Artesunate treatment. Nevertheless, consistent with our hypothesis that Erk1/2 activated DAPK1 (Supplementary Figure 4C), Artesunate treatment alone resulted in lamellipodia loss (Figure 5C), reminiscent of DAPK1 dephosphorylation [18]. This phenomenon was reversed in the presence of U0126 in HepG2 cells treated with Artesunate (Figure 5C). Furthermore, Artesunate treatment elevated Caspase-3 at 6 h post-treatment (Supplementary Figure 4D). Caspase activity was attenuated by U0126 addition after 12 h post-treatment by CaspaseGlo^®^ assay (Figure 5D) and Sensor C3 reporter assay (Supplementary Figures 4E and 4F), respectively. It was noteworthy that U0126 did not eventually rescue HepG2 cells from apoptosis during Artesunate treatment (Figure 5D). Instead, U0126 treatment alone reduced cell numbers after 36 h post-treatment as compared to control (Figure 5D). Because the starting cell density reached 10,000 cells/well on 96-well plate to obtain sufficient CaspaseGlo^®^ signal, we next tested cell proliferation using a starting cell density of 5,000 cells/well on 96-well plate.

Collectively, these data supported our hypothesis that Artesunate induced Erk1/2 phosphorylation and subsequent DAPK1 dephosphorylation to promote apoptosis *in vitro*.

**Figure 5.**
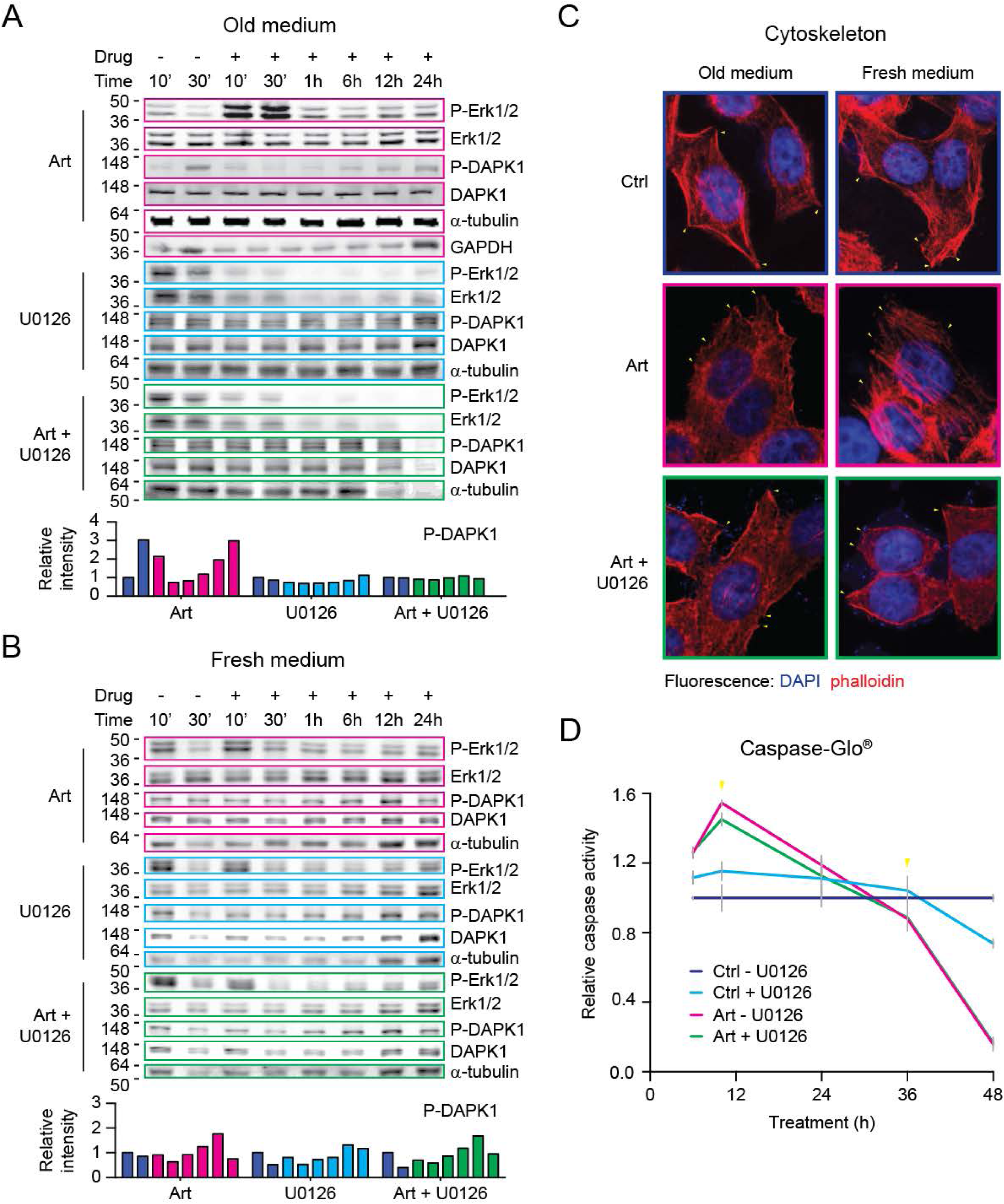
Artesunate inhibited HepG2 cell proliferation via Erk1/2-DAPK1 pathway *in vitro*. (A) Western blotting of Erk1/2-DAPK1 pathway proteins after direct Artesunate addition. Western blotting was performed on HepG2 cells treated with 30 μM Artesunate and/or 5 μM U0126 by direct drug addition (old medium) over a time course of 24 h. Parallel blotting of phosphorylated proteins was performed; α-tubulin and GAPDH was used as loading controls. The graph denoted quantification results of phosphorylated DAPK1 bands in corresponding Western blots. (B) Western blotting of Erk1/2-DAPK1 pathway proteins after Artesunate addition by medium replacement. Western blotting was performed on HepG2 cells treated with 30 μM Artesunate and/or 5 μM U0126 by medium replacement (fresh medium) over a time course of 24 h. Parallel blotting of phosphorylated proteins was performed; α-tubulin was used as loading control. The graph denoted quantification results of phosphorylated DAPK1 bands in corresponding Western blots. (C) Actin staining of HepG2 cells treated with Artesunate by different experimental procedures. HepG2 cells were treated with 30 μM Artesunate in the absence and presence of 5 μM U0126 by direct drug addition (old medium) or medium replacement (fresh medium), and subject to IF staining with DAPI (nucleus), α-tubulin antibody (channel not shown for clarity) and phalloidin (β-actin), followed by fluorescence imaging under 40x objective. Yellow arrows indicated lamellipodia. (D) Caspase activation assay on Artesunate-treated HepG2 cells. Caspase-Glo^®^ assay was performed on HepG2 cells treated with 30 μM Artesunate and/or 5 μM U0126 over a time course of 48 h. The mean and standard deviation from quadruplicate experiments of relative caspase activity were plotted against time (h). Yellow arrows indicated critical time point(s).

### U0126 did not rescue HepG2 cells from Artesunate treatment because U0126 triggered cell multiploidy independent of Artesunate treatment

Firstly, Alamar Blue staining showed that U0126 addition stimulated cell growth in HepG2 cells during double treatment with DMSO or Artesunate before 48 h post-treatment (Figure 6A). However, although cell proliferation rate remained high in U0126-treated HepG2 cells, proliferation slowed down in cells treated with Artesunate and U0126 after 48 h incubation (Figure 6A). Cell proliferation eventually leveled off in double treated cells to reach the same level as compared to Artesunate treatment alone at 72 h post-treatment (Figure 6A). In contrast, U0126 treatment alone led to higher proliferation than control cells until 72 h post-treatment (Figure 6A).

**Figure 6.**
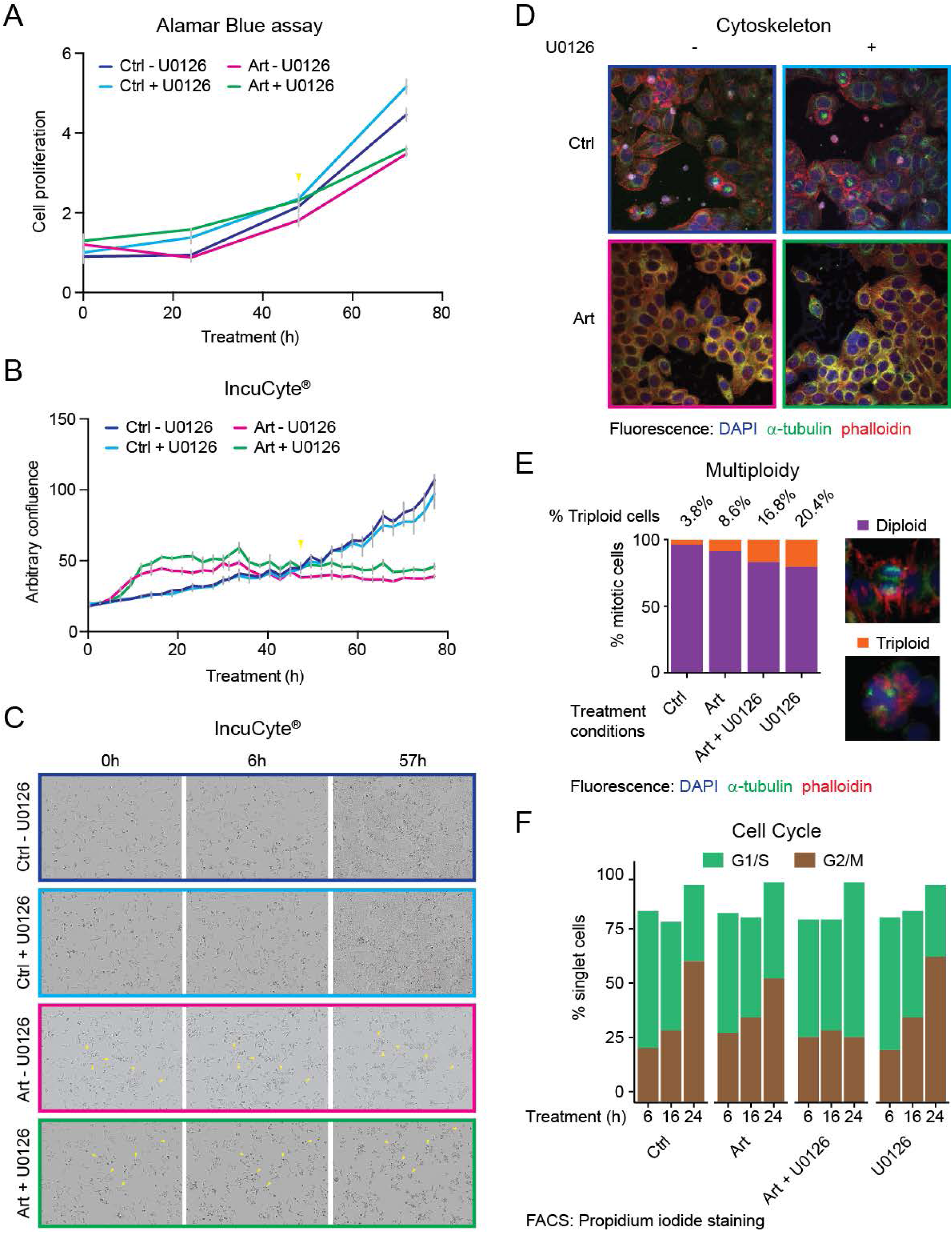
Multiploidy formation induced by U0126 abrogated its rescue effect on Artesunate-treated HepG2 cells. (A) Cell proliferation assay of HepG2 cells under Artesunate treatment alone or in combination with U0126. Alamar Blue assay was performed on HepG2 cells treated with 30 μM Artesunate and/or 5 μM U0126 over a time course of 72 h. The mean and standard deviation of normalized cell viability was plotted against time. Yellow arrows indicated critical time point(s). (B) Cell proliferation assay of HepG2 cells under Artesunate treatment alone or in combination with U0126. IncuCyte^®^ time-lapse imaging was performed on HepG2 cells treated with 30 μM Artesunate and/or 5 μM U0126 over a time course of 74 h. The mean and standard deviation of arbitrary confluence from 4 field of views in triplicate experiments were plotted against time (h). Yellow arrows indicated critical time point(s). (C) Cell morphology analysis of IncuCyte^®^ time-lapse images. Images of the IncuCyte^®^ time-lapse imaging experiment was analyzed to observe changes in cell morphology in HepG2 cells treated with 30 μM Artesunate and/or 5 μM U0126 over a time course of 74 h. Yellow arrows indicated critical morphological changes. (D) IF staining of cytoskeleton of HepG2 cells under Artesunate treatment alone or in combination with U0126. HepG2 cells treated with 30 μM Artesunate and/or 5 μM U0126 for 12 h were fixed and stained against DAPI (nucleus), α-tubulin antibody and phalloidin (β-actin), followed by fluorescence imaging under 20x objective. (E) Multiploidy analysis of HepG2 cells under Artesunate treatment alone or in combination with U0126. Analysis of the IF staining result of HepG2 cells treated with 30 μM Artesunate and/or 5 μM U0126 for 12 h was conducted to count the number of diploid and triploid cells among centrosome-distinguishable mitotic cells. (F) Cell cycle analysis of HepG2 cells under Artesunate treatment alone or in combination with U0126. Propidium iodide staining of fixed HepG2 cells treated with 30 μM Artesunate and/or 5 μM U0126 over a time course of 24 h was analyzed by fluorescence activated cell sorting.

To investigate the growth defect of Artesunate and U0126 double treated cells, we carried out IncuCyte^®^ *in situ* time-lapse imaging and confluence measurement of HepG2 cells under single or double treatment with 30 μM Artesunate and 5 μM U0126 respectively. IncuCyte^®^ analysis showed that HepG2 cells treated with Artesunate alone or in combination with U0126 rapidly reached 50% confluence within 10 h, but cell growth retarded until 74 h post-treatment (Figure 6B). Cell confluence was slightly higher in Artesunate and U0126 double treated cells as compared to Artesunate treatment alone (Figure 6B). Alternatively, DMSO-treated cells in the absence or presence of U0126 continuously proliferate to reach 100% confluence at 74 h post-treatment without significant difference between the two conditions (Figure 6B). Examination of the cell morphology of HepG2 cells prevailed no difference between cells treated with DMSO and U0126 (Figure 6C). However, Artesunate-treated cells lost suppleness, while lamellipodia were present (Figure 6C). In contrast, most of the Artesunate and U0126 double treated cells became round after 48 h post-treatment (Figure 6C).

Although proliferation curves differed between measurements made by Alamar Blue assay (Figure 6A) and IncuCyte^®^ imaging (Figure 6B) at early time points, proliferation of HepG2 cells treated with either U0126 or DMSO overtook the Artesunate-treated and double treated cells at 48 h post-treatment in both assays, indicating that this time point was critical to loss of rescue by U0126. During morphological analysis, we observed multiploid formation in U0126-treated HepG2 cells independent of Artesunate treatment (Figure 6D). Examination of over 6,000 centrosome-distinguishable mitotic cells under IF staining for each treatment condition showed that 96.2% of DMSO-treated HepG2 cells underwent diploid mitosis, whereas U0126 resulted in formation of triploids in 20.4% of cells and, to a much lesser extent, multiploids (Figure 6E). In parallel, double treatment with Artesunate and U0126 resulted in 16.8% of triploids (Figure 6E). Artesunate-treated cells showed 8.6% of triploids (Figure 6E), suggesting that no accumulative effect occurred during double treatment.

Next, cell cycle analysis by fluorescence activated cell sorting (FACS) demonstrated that the percentage of G2/M cells under DMSO treatment gradually increased from 24.8%, 33.1% to 65.2% at 6 h, 16 h and 24 h post-treatment, respectively (Figure 6F). The percentage of G2/M cells also increased over time in HepG2 cells treated with Artesunate (31.8%, 39.0% and 56.5%) or U0126 (24.1%, 39.3% and 66.8%) for 6 h, 16 h and 24 h post-treatment, respectively (Figure 6F). However, cell cycle arrest in G1/S phase was observed in HepG2 cells under Artesunate and U0126 double treatment starting at 16 h post-treatment, where the percentage of G2/M cells leveled off from 30.2% to 33.2% at 6 h and 16 h post-treatment, then fell to 29.5% at 24 h post-treatment, respectively (Figure 6F). Analysis of the DNA content in FACS showed a tiny peak after the G2/M peak in HepG2 cells treated with U0126 (6.8%) and Artesunate and U0126 (5.1%), but not in control (2.1%) nor Artesunate-treated cells (3.0%) (Supplementary Figure 5A), suggesting of multiploidy emergence in U0126-treated cells independent of Artesunate treatment.

The evasion of cell cycle arrest in HepG2 cells under U0126 treatment alone in comparison to double treatment with Artesunate could be explained by the continuous cell turnover in HepG2 cells regardless of insult, as observed in persistent Caspase-3 cleavage (Supplementary Figure 4D). We hypothesized that the continuous cell turnover assisted HepG2 cells under U0126 single treatment to remove multiploids, so as to remove cells under proliferative adversity. However, in the presence of Artesunate, multiploids independently caused by U0126 could not be efficiently removed, resulting in cell cycle arrest at 24 h post-treatment (Figure 6F). This eventually led to retarded cell proliferation and cell death in double treated cells, seen as the turning point at 48 h post-treatment in cell proliferation assays (Figure 6A and 6B). Collectively, these data provided evidence for the failure of U0126 to rescue HepG2 cells from death under Artesunate treatment.

Taken together, our results demonstrated that U0126 inhibition of MEK1-Erk1/2 pathway led to early proliferation advantage in HepG2 cells under Artesunate treatment, but its virtue was eventually overcome by cell multiploidy followed by cell cycle arrest.

## Discussion

In this study, we used Artesuante-treated HepG2 cells as a model to investigate the variance between *in vitro* and *in vivo* drug screening. For the *in vitro* models, monolayer culture was used for our initial drug screen; alternatively, organoid culture was used for biomarker staining because organoids partially mimicked the three-dimensional tissue architecture *in vivo*. On the other hand, we constructed the *in vivo* models of subcutaneous and liver implantation xenograft mice, respectively. The subcutaneous xenograft model was assumed to possess normal liver function, whereas the liver implantation model simulated real-life clinical situation. These models have all been used in preclinical screening of anti-cancer drugs. In this study, we applied Artesunate treatment of HepG2 cells to these models to decipher the discrepancy between *in vitro* and *in vivo* screening. Our results depicted that: (1) Artesunate was most effective in inhibiting monolayer HepG2 cells, followed by HepG2 organoids, and was ineffective in our subcutaneous xenograft model; (2) AFP depicted tissue architecture-dependent expression, but was neither a biomarker nor response indicator of effective Artesunate treatment *in vitro*; (3) albumin was a response indicator *in vitro* but not *in vivo* (Figure 7A).

**Figure 7.**
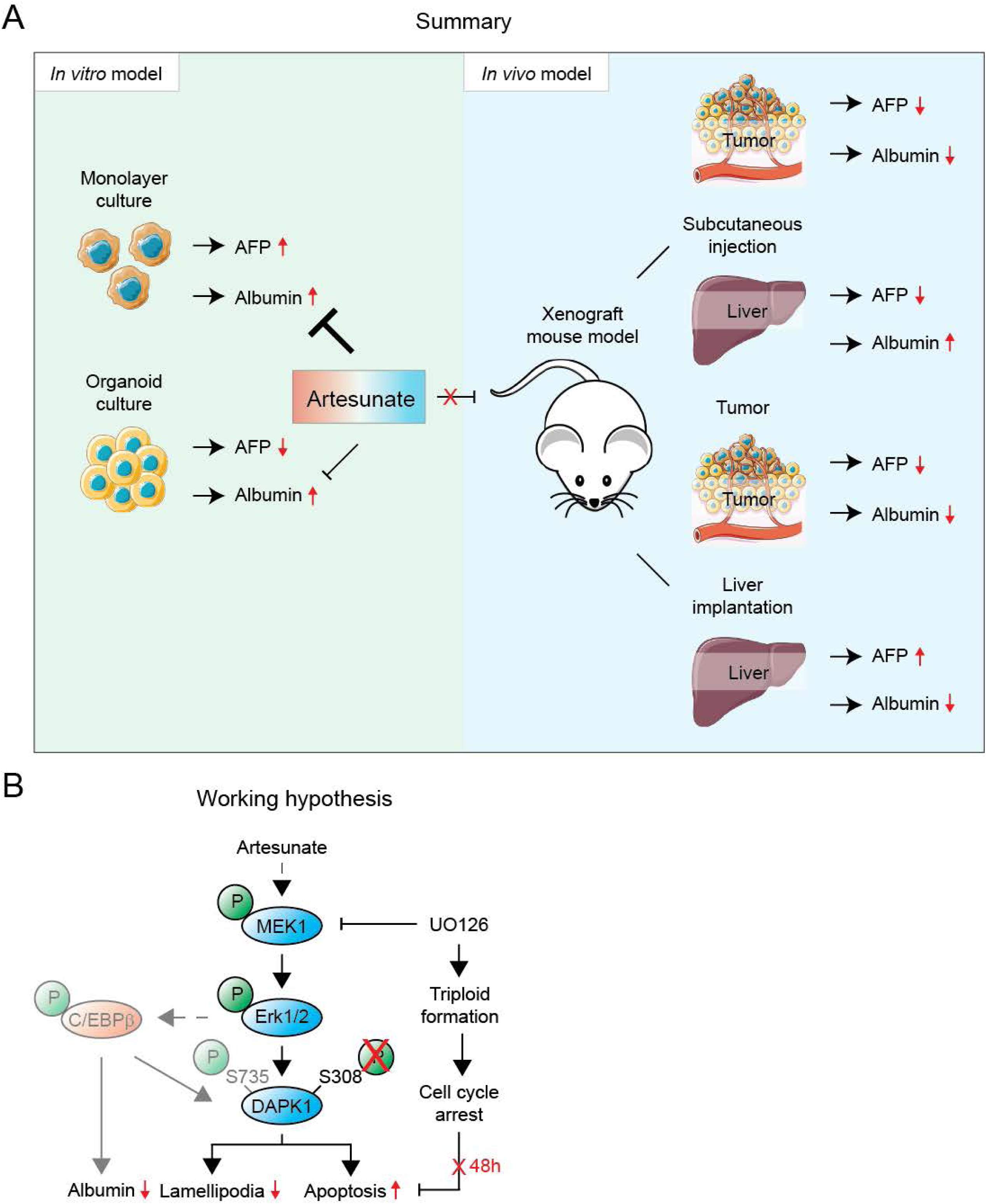
Albumin was a response indicator *in vitro*. (A) Comparison between *in vitro* and *in vivo* models. AFP depicted tissue architecture-dependent expression *in vitro* and was neither a biomarker nor response indicator in our comparison model. Alternatively, albumin was a response indicator *in vitro*. (B) Working hypothesis of Artesunate effects *in vitro*. Artesunate triggered lamellipodia loss and apoptosis via Erk1/2-DAPK1 pathway in HepG2 cells *in vitro*, which was not reversed by U0126 due to its independent effect of triggering multiploid formation. Greyed out portions of the hypothesis were not supported by empirical evidence in this study.

Next, we showed that Artesunate inhibited HepG2 cell proliferation and activated the Erk1/2-DAPK1 pathway to promote apoptosis *in vitro* (Figure 7B). In concert to our results, strong Erk2 phosphorylation was induced by hepatocyte growth factor (HGF), leading to retarded cell proliferation in HepG2 cells, which was reversed by adding the MEK inhibitors of PD98059 or U0126 that alleviated Erk2 phosphorylation [24]. Similarly, activation of Erk1/2-DAPK1 pathway has been reported to trigger apoptosis in NIH3T3 cells under starvation [18]. Contrariwise, Artemisinin was shown to protect human retinal pigment epithelial cells through Erk/CREB signaling [25], and conferred neuroprotection to PC12 cells *in vitro* [26]. Hence, the consequence of Erk phosphorylation might be context-dependent.

More interestingly, our Western blots showed different Erk phosphorylation patterns during direct drug addition and medium replacement, suggesting that experimental conditions could interfere interpretation. Notwithstanding, DAPK1 dephosphorylation and lamellipodium loss was observed in Artesunate-treated cells but not in control in both scenarios, indicating that the downstream pathways were conserved. It was notable that no commercial antibody against DAPK1-pSer735 was available, so we tested the inhibitory phosphorylation site on Ser308, which was reported to be crucial for its kinase activity [27, 28]. Hence, DAPK1 dephosphorylation observed in our experiments indicated DAPK1 activation.

Intracellular albumin reduction appeared to be a response indicator of effective Artesunate treatment *in vitro*. Using genomic sequence alignment, we identified a downstream Erk target that bound to the albumin promoter, named CCAAAT/enhancer-binding protein-beta (C/EBPβ), by TFmapper (http://www.tfmapper.org/). Overexpression of C/EBPβ was reported to downregulate albumin transcription in HepG2 cells *in vitro* [29]. Because albumin was reported to promote proliferation triggered by Erk activation in kidney cells [30, 31], reducing albumin may hinder Erk-mediated cell proliferation. Coincidently, C/EBPβ upregulated DAPK1 expression and induced apoptosis in mouse embryonic fibroblasts upon interferon-gamma (IFNγ) stimulus [32]. Therefore, C/EBPβ might provide parallel pathways to impede growth and promote death in HepG2 cells under Artesunate treatment in our current model (Figure 7B). Nevertheless, in lieu of the modest effects of U0126 on caspase activation and cell proliferation in Artesunate-treated HepG2 cells before 24 h post-treatment, we could not rule out the possibility of co-stimulatory pathway(s) in addition to Erk signaling, which required further investigation.

Given that tissue architecture did not confer to drug response and no changes on animal physiology was perceived, we wondered if the pharmacokinetics of Artesunate might attribute to drug inefficacy *in vivo*. Indeed, Morris, *et al*, demonstrated that the blood clearance rate and half-life of Artesunate was 2-3 L kg^-1^ h^-1^ and less than 15 min for intravenous injection in human patients [33]. The clearance rate and half-life was slightly higher, 19.2-30 L kg^-1^ h^-1^ and 20-45 min for oral administration, and 2.4-3.5 L kg^-1^ h^-1^ and 25-48 min for intramuscular administration, respectively [33]. Therefore, we reasoned that mice would exhibit similar clearance rate *in vivo*, thus resulting in no drug response due to rapid drug clearance. Nevertheless, given its tumor inhibition efficiency *in vitro* and mild effects *in vivo*, we hypothesized that improving drug distribution might improve Artesunate efficacy *in vivo*.

## Materials and Methods

### Drugs and Reagents

Artesunate (CAS# 88495-63-0) and Dihydroartemisinin (CAS# 71939-50-9) were bought from Shanghai Aladdin Bio-Chem Technology Company Limited, China. Artemisinin (CAS# 63968-64-9) and Artemether (CAS# 71963-77-4) were bought from Carbosynth Limited, UK. Cell culture medium and auxiliary components were bought from Life Technologies, USA. Matrigel^®^ (Cat# 356235) was bought from Corning, USA. All other reagents were bought from Sigma-Aldrich, USA, unless otherwise indicated. Antibodies used in this study were listed in Supplementary Table 1.

### Cell culture

The indicated cell lines were obtained from Professor Chuxia Deng’s laboratory. Most cell lines were passaged 1-2 times before experiments and frozen immediately after use. 1 mL culture medium was taken from the first passage and sent to the Bioimaging Core of University of Macau, for mycoplasma testing by MycoAlert^TM^ Mycoplasma Detection Kit (Lonza, Switzerland) following manufacturer’s protocol.

All cells were cultured on monolayer with Dulbecco’s Modified Eagle Medium (DMEM) or RPMI 1640 medium supplemented with 10% fetal bovine serum (FBS), 100 U/mL Penicillin-Stretomycin and 2 mM L-glutamine. Cells were incubated at 37°C in a 5% CO_2_ humidified incubator and passaged every 3-4 d.

For HepG2 organoid culture, HepG2 cells passaged on monolayer were resuspended as 30,000 cells/mL final concentration in 1:1 (v/v) Matrigel^®^ : 3D medium comprising of Advanced DMEM/F12 medium supplemented with 100 U/mL Penicillin-Stretomycin, 1% (v/v) GlutaMAX^TM^, 10 mM HEPES pH7.4, 1x B-27^TM^ Supplement without Vitamin A, 1x N-2 supplement, 1.25 mM N-acetyl-L-cysteine, 10 mM Nicotinamide, 10 nM recombinant human [Leu^15^]-Gastrin I, 50 ng/mL recombinant human Epidermal Growth Factor (EGF), 10 μM Forskolin, 5 μM A8301 (Selleckchem, USA) and 10 μM Y27632 (Selleckchem, USA). Resuspended cells were seeded in 30uL droplets at the bottom of 6-well plate and 2 mL 3D medium was applied. Cells were incubated at 37°C in a 5% CO_2_ humidified incubator and passaged every 4 d.

For immunofluorescence staining, organoids were cultured in 3D medium supplemented with 1% (w/v) of 1:2 (w/w) gelatin : agar on 4-well chamber slides, and subject to fixation.

### Drug screening and IncuCyte^®^ analysis

Cells were seeded at 2,000-8,000 cells/well in 30 μL cell culture medium on a 384-well clear-bottom square-well plate. 3 μL of serially diluted 10x concentration of each drug, yielding a dose gradient from 200 μM to 12.8 nM final concentration, was added to each well in quadruplicates immediately after cell seeding. The cells were incubated at 37°C in a 5% CO_2_ humidified incubator for 72 h. Relative cell viability was measured by fluorescence at ex. 560 nm / em. 590 nm using Alamar Blue assay [34] on Molecular Devices SpectraMax M5 Plate Reader. 0.1% dimethyl sulfoxide (DMSO) was used as negative control and cell-free medium was setup as blank control.

For IncuCyte^®^ analysis, HepG2 cells were seeded at 5,000 cells/well in 100 μL cell culture medium on a 96-well plate. The cells were incubated at 37°C in a 5% CO_2_ humidified incubator overnight before a final concentration of 30 μM Arteusnate and/or 5 μM U0126 was applied. Immediately after drug addition, the cells were transferred to Satorius IncuCyte^®^ Live Cell Imaging System. Each treatment condition contained triplicate wells, where 4 field of views (FOVs) per well were imaged at 2 h intervals for 74 h in total. 0.1% DMSO was used as negative control.

### Intracellular iron measurement

Sampled tissue from different mouse organs were freshly cut immediately after mouse euthanization, followed by three washes with phosphate buffered saline (PBS). Sampled cells were grown to confluence in a 6-well plate under normal culture conditions, trypsinized, counted and washed with PBS thrice. Afterwards, the samples were freeze-dried into dry powder and weighed. 1 mL concentrated nitric acid was applied to dissolve all samples by microwave. The extract was diluted and metered to 50 mL with pure water. 5 mL of the diluted extract was sent for ionization coupled plasma atomic emission spectroscopy (ICP-AES) analysis at the Analytic Centre of Jinan University, Guangzhou, China. The measured iron concentrations were finally converted to mg/g tissue (dry weight) or pg/cell based on arithmetic calculation.

### Cell cycle analysis

HepG2 cells were initially treated with 0.1% DMSO or 10-50 μM Artemisinin and its derivatives on a 6-well plate at 37°C in a 5% CO_2_ humidified incubator for 12 h. Cells were trypsinized and transferred to 4 mL round-bottom tubes. 2 volumes of pure ethanol were added in a dropwise manner on slow vortex. After the cells were fixed, final concentration of 5 μg/mL propidium iodide (PI) (Life Technologies, USA) and 1 mg/mL RNase I (Life Technologies, USA) was added to stain the cells for 10 min. Lastly, the cells were filtered through 40 μm cell strainer and loaded on BD FACS Calibur Flow Cytometer for cell cycle analysis by flow cytometry.

### Radical oxygen species (ROS) staining

HepG2 cells were treated with doses of 10-50 μM Artemisinin and its derivatives on a 6-well plate at 37°C in a 5% CO_2_ humidified incubator for 12 h. After drug treatment, cells were first washed with PBS thrice. Then a final concentration of 10 μM Carboxyl-H_2_DCFDA (Life Technologies, USA) was added to each well and incubated at room temperature for 10-20 min. Cells were immediately imaged under green fluorescence channel at 10x magnification using Life Technologies EVOS FL Imaging System. Carbonylcyanide-4-(trifluoromethoxy)-phenylhydrazone (FCCP) (Enzo Life Sciences, Hong Kong) was used as positive control, 0.1% DMSO was used as negative control.

### Intracellular pH measurement

HepG2 cells were initially treated with 0.1% DMSO or 30 μM of Artesunate on a 24-well plate at 37°C in a 5% CO_2_ humidified incubator for 12 h. Then a final concentration of 0.5 μM 2’,7’-bis-(2-carboxyethyl)-5-(and-6)-carboxyfluorescein (BCECF) (Life Technologies, USA) was added to each well and incubated at room temperature for 30 min, followed by plate reading at ex. 440 nm / em. 535 nm and ex. 490 nm / em. 535 nm using Molecular Devices SpectraMax M5 Plate Reader. A standard curve was plotted using home-made buffers constituting a pH gradient from 6.5 to 8.5.

### Western blotting

HepG2 cells were initially treated with 0.1% DMSO or 30 μM of Artesunate, in the absence and presence of 5 μM U0126, on a 6-well plate at 37°C in a 5% CO_2_ humidified incubator for the indicated time on each figure. Cells were then washed with PBS thrice. Subsequently, gel loading dye was directly applied to the adherent cells, incubated at room temperature for 10 min, and transferred to heat-block for protein denaturation at 95°C for 10 min. Protein samples were separated on 10% or 15% SDS-PAGE followed by Western blotting using specific antibodies. Blots were scanned by near-infrared fluorescence using Licor Odyssey CLx Imager.

### Immunofluorescence

HepG2 cells in monolayer or organoid culture were initially treated with 0.1% DMSO or 30 μM of Artesunate, in the absence and presence of 5 μM U0126, on a 4-well chamber slide at 37°C in a 5% CO_2_ humidified incubator for 12 h. Cells were then washed with PBS thrice. Subsequently, cells were fixed with 10% Formalin and washed with 1% Triton X-100, followed by specific antibody staining. 10x to 40x magnified images were taken using Olympus BX83 Upright Fluorescent Microscope, whereas 63x magnified images were taken on Carl Zeiss LSM 710 Confocal Fluorescent Microscope.

### Xenograft mouse model

All mouse experiments were conducted in the Animal Facility of University of Macau, after approval by the Animal Research Ethics Committee of University of Macau under protocol UMARE-050-2017.

For the subcutaneous xenograft mouse model, 3.0 × 10^6^ HepG2 cells suspended in 1:1 (v/v) Matrigel^®^ : PBS was implanted subcutaneously on the left and right flank of athymic nude mice (Balb/c Nude). Tumorigenesis occurred between 10-14 d. When tumors reached 2 mm in any dimension, the mice were separated into two groups of 10 mice. Each group of mice was subject to intraperitoneal injection of 50 mg/kg Artesunate in 0.4% Tween-80 in PBS and solvent alone, respectively, at a frequency of 2 times per week for a total span of 4 weeks or until the tumor exceeded 1.5 cm^3^ in any dimension.

For the liver implantation xenograft mouse model, the liver of athymic nude mice (Balb/c Nude) was first exposed through ventral midline incision after anesthesia by 500 mg/kg Avertin. Sensor C3-labelled HepG2 cells were suspended in PBS at a final concentration of 4 × 10^7^ cells/mL. 50 μL of the cell suspension was slowly injected using a 29G needle inserted 5 mm into the left and right anterior lobe of the liver respectively, until the liver turned pale. After surgery, the abdomen was closed in two layers using surgical thread. Mice were kept on a 37°C hot plate until revival and returned to their cages consequently. Tumors were allowed to grow for 8 weeks, followed by tumor resection after euthanasia.

Before euthanasia, mice underwent anesthesia by 250 mg/kg Avertin. Blood was extracted from the heart with ethylenediaminetetraacetic acid (EDTA) as anti-coagulant. Tumors and other relevant organs were resected after euthanasia.

### Hematoxylin and eosin (H&E) staining and immunohistochemistry (IHC)

Resected tumors and organs were immersed in Thermo Fisher Cryochrome™ embedding resin, and stored at −80°C until use. The frozen blocks were cut into 5-10 μm slices using Leica CM3050S Cryostat. The slices were transferred to 25 × 75 mm positive adhesion glass slides and fixed in 10% Formalin, followed by H&E staining following standard procedures or antibody staining following manufacturer’s instructions for Thermo Fisher Histostain-*Plus* IHC Kit. IHC slides were counterstained by Hematoxylin. All slides were dehydrated by xylene and mounted by DPX Mountant. 10x magnified images were taken using Olympus BX83 Upright Fluorescent Microscope.

### Blood biochemistry

Immediately after extraction, whole blood was centrifuged at 10,000g at room temperature for 2 min. The supernatant was aspirated and transferred to a new microfuge tube for storage at −80°C if necessary. Blood serum was loaded on Sysmex BX-3000 Automated Chemistry Analyzer calibrated using Biosystems Biochemistry Multi-calibrator and Biosystems Lipid Control Serum II, respectively. Results were automatically calculated by the complimentary software.

### Isothermal calorimetry

Isothermal calorimetry (ITC) was performed in the Drug Development Core, Faculty of Health Sciences, University of Macau. 200 μM Artesunate was titrated into 5μM bovine serum albumin (BSA) at 25°C on GE Healthcare Microcal iTC200 system. The solvent was phosphate buffered saline (PBS); BSA solvent was supplemented with 0.1% DMSO to match the ligand solvent.

## Authors’ Contribution

S.L.S., A.H.W. and C.D. conceived this project. J.K.W., S.L.S. and K.H.W. were supervised by A.H.W. A.H.W. designed experiments and analyzed results. J.K.W., S.L.S., K.H.W. and A.H.W. performed experiments; H.W. wrote R scripts for data analysis; F.X. constructed the Sensor C3-labelled HepG2 cell line; Y.Q. provided 3D medium and instructions on organoid culture; M.Z. and L.Z. performed liver implantation surgeries. All authors contributed to preparation of this manuscript.

## Acknowledgement

Acknowledgements to Mr. Zepeng Jiao of Analytic Centre, Jinan University for instructions on ICP-AES, Dr. Peng Lyu of Prof. Henry Kwok’s lab for instructions on IncuCyte^®^, Dr. Qiang Chen of Prof. Chuxia Deng’s lab and Dr. Weiwei Liu of Bioimaging Core for supervision on FACS, Dr. Xiaohui Hu of Drug Development Core for supervision and assistance with ITC, Mr. Joshua Cai, Ms. Avery Tam and Mr. Kevin Tang of Animal Facility for supervision and assistance on mouse experiments, Dr. Jammy Chan of Kiang Wu Hospital for advice on histology, Prof. Wenhua Zheng for provision of Artemisinin and analogs, Prof. Qian Luo for provision of Sensor C3 reporter plasmid, and Prof. Joong Sup Shim for provision of BCECF. We specially thank our former colleague Prof. Ruihong Wang for rigorous comments on this manuscript. All authors read and approved of publication of this manuscript.

