## Supplementary Figures for "Dissecting the dilemma between *in vitro* and *in vivo* drug screening: treating HepG2 cells with Artesunate as a model"

#### Supplementary Figure Legends

**Supplementary Figure 1.** Staining of HCC biomarkers in parallel untreated nude mice. (A) IHC staining of AFP in untreated nude mice in parallel to the corresponding xenograft models. Livers and kidneys of untreated nude mice were fixed and stained with AFP antibody. Brightfield images were taken using 4x objective. (B) IHC staining of albumin in untreated nude mice in parallel to the corresponding xenograft models. Livers and kidneys of untreated nude mice were fixed and stained with albumin antibody. Brightfield images were taken using 4x objective.

**Supplementary Figure 2.** Artesunate did not bind to albumin. (A) Molecular docking of Artesunate on human blood serum albumin. Molecular docking of Artesunate (CAS number: 88495-63-0) on human albumin protein (PDB ID: 1N5U) was performed by Swiss Dock (http://www.swissdock.ch/). (B) Isothermal calorimetry of Artesunate against bovine serum albumin. 100 μM Artesunate (CAS number: 88495-63-0) was titrated against 5 μM bovine serum albumin in PBS buffer supplemented with 0.1% DMSO. Graph was plotted by Origin Lab software supplied with the instrument. (C) Sequence alignment of albumin proteins of different species. Peptide sequences of albumin proteins from human (Uniprot ID: P02768), bovine (Uniprot ID: P02769) and mouse (Uniprot ID: P07724) was aligned on Uniprot website (https://www.uniprot.org) by Clustal Omega. Colored triangular annotations were made in accordance to predicted binding sites of Artesunate on human albumin protein by Swiss Dock.

**Supplementary Figure 3.** Artesunate did not inhibit HepG2 cell proliferation by known Artemisinin modes of action. (A) Iron measurement of mouse organs. Organs were dissected from one untreated athymic nude mouse after euthanasia, and subject to ionization coupled plasma atomic emission spectroscopy for iron measurement. Iron concentration per gram (mg/g) of tissue was plotted against the investigated tissue. (B) Correlation between $IC_{50}$ values after Artesunate treatment and intracellular iron concentration of all investigated cell lines. $IC_{50}$ values of the eleven investigated cell lines after Artesunate treatment were plotted against corresponding intracellular iron concentration of respective cell lines. Each dot represented one cell line and the color of the dot denoted the tissue origin of the indicated cell line. (C) FACS of Artesunate-treated HepG2 cells. 30 μM Artesunate was applied to HepG2 cells and treated for 12 h, and subject to fluorescence activated cell sorting after propidium iodide staining. (D) ROS staining of Artesunate-treated HepG2 cells. HepG2 cells were treated with $IC_{50}$ concentrations of the four Artemisinin analogs for 12 h, and subject to carboxy-$H_2$DCFDA staining followed by fluorescence imaging under 10x objective for radical oxygen species detection. FCCP and DMSO were used as positive and negative controls respectively. (E) pH measurement of Artesunate-treated HepG2 cells. HepG2 cells were treated with $IC_{50}$ concentrations of the four Artemisinin analogs for 12 h, and subject to BCECF staining followed by fluorescence plate reading for pH measurement. Each dot represented one data point, whereas the mean and standard deviation was represented by the column and error bar. Standard curve measurements from standard pH buffers was used for pH calibration.

**Supplementary Figure 4.** Analysis of the cell death mechanism of Artesunate-treated HepG2 cells. (A) Western blotting of Artesunate-treated HepG2 cells against different kinase antibodies. 30 $\mu$M Artesunate was applied to HepG2 cells and treated over a time course of 24 h. Parallel blotting of phosphorylated and total proteins was performed. (B) Cell death assay on Artesunate-treated HepG2 cells. IncuCyte® time-lapse imaging was performed on HepG2 cells treated with 30 $\mu$M Artesunate and/or 5 $\mu$M U0126 over a time course of 30 h. The mean and standard deviation of arbitrary confluence from 4 field of views in triplicate experiments were plotted against time (h). (C) Gene structure of DAPK1 kinase. DAPK1 is a multi-domain kinase that participates in apoptosis and autophagy. (D) Western blotting of cell death-related proteins in Artesunate-treated HepG2 cells. 30 $\mu$M Artesunate was applied to HepG2 cells and treated over a time course of 24 h. (E) Sensor C3 measurement of Artesunate-treated HepG2 cells with or without U0126. Sensor C3-labelled HepG2 cells were treated with 30 $\mu$M Artesunate in the absence or presence of 5 $\mu$M U0126 over a time course of 24 h, and subject to time-lapse fluorescence imaging under 10x objective for FRET detection. The mean and standard deviation of the FRET ratio of 4-6 field of views were plotted against time (h) starting from drug addition. (F) Real-time analysis of Sensor C3-labelled HepG2 cells under Artesunate treatment with or without U0126. Analysis of real-time images of Sensor C3-labelled HepG2 cells under Artesunate treatment with or without U0126 was conducted to look for temporal patterns of Caspase-3 activation. Each column denoted each field of view, whereas the color intensity indicated mean FRET ratio in each field of view at the indicated time point.

**Supplementary Figure 5.** (A) Cell cycle analysis of multiploid formation triggered by U0126. Propidium iodide staining of fixed HepG2 cells treated with 30 $\mu$M Artesunate and/or 5 $\mu$M U0126 for 16 h was analyzed by fluorescence activated cell sorting.

### Supplementary Figure 1

A

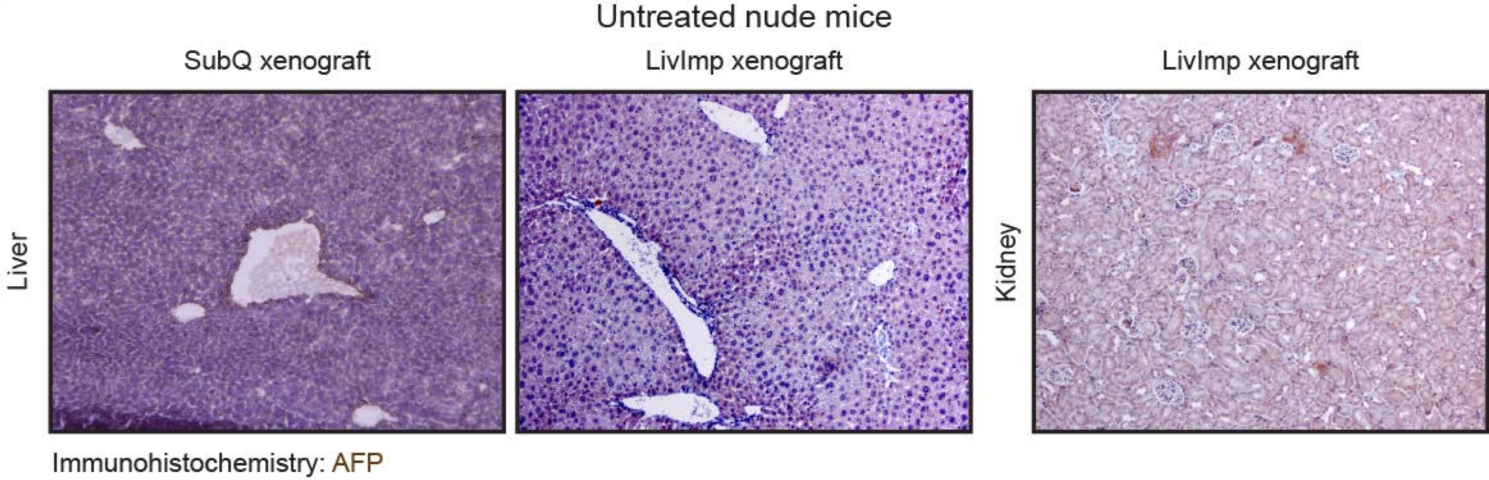


B

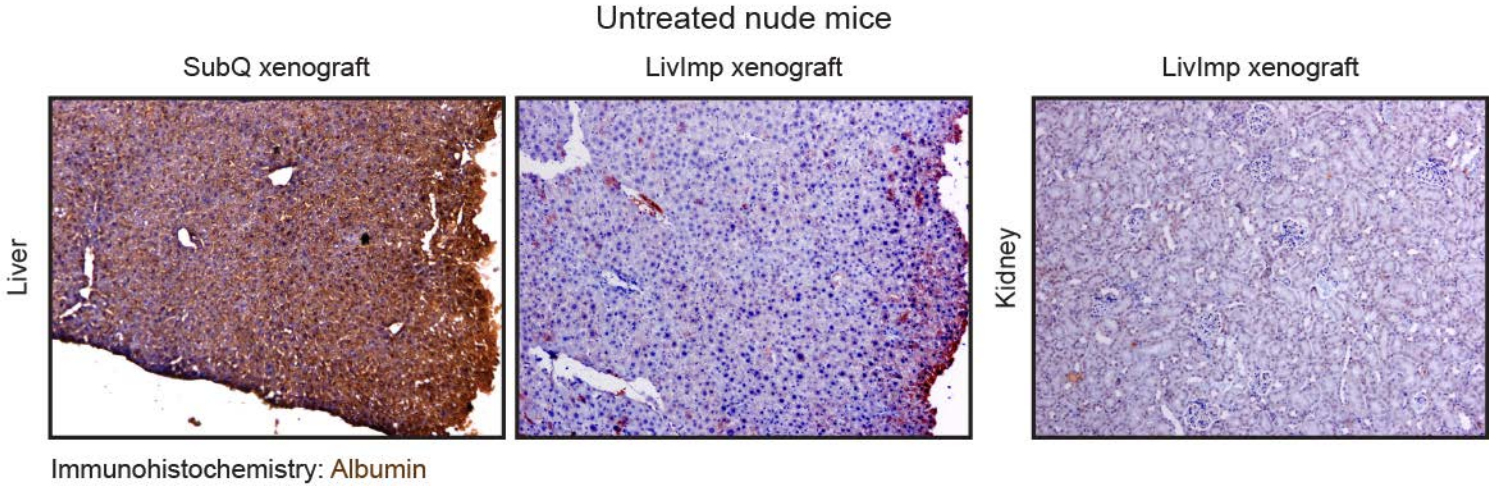

### Supplementary Figure 2

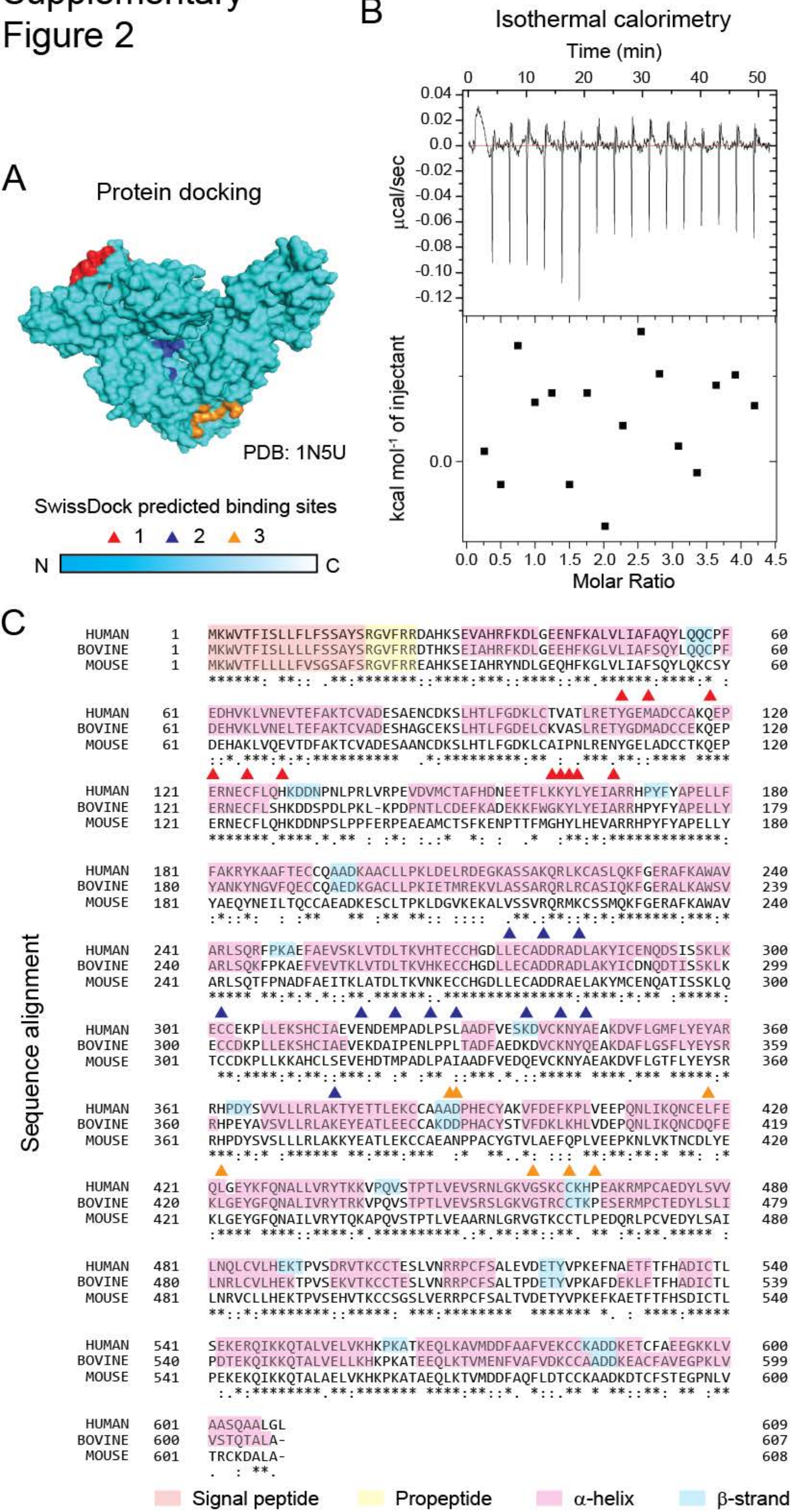

### Supplementary Figure 3

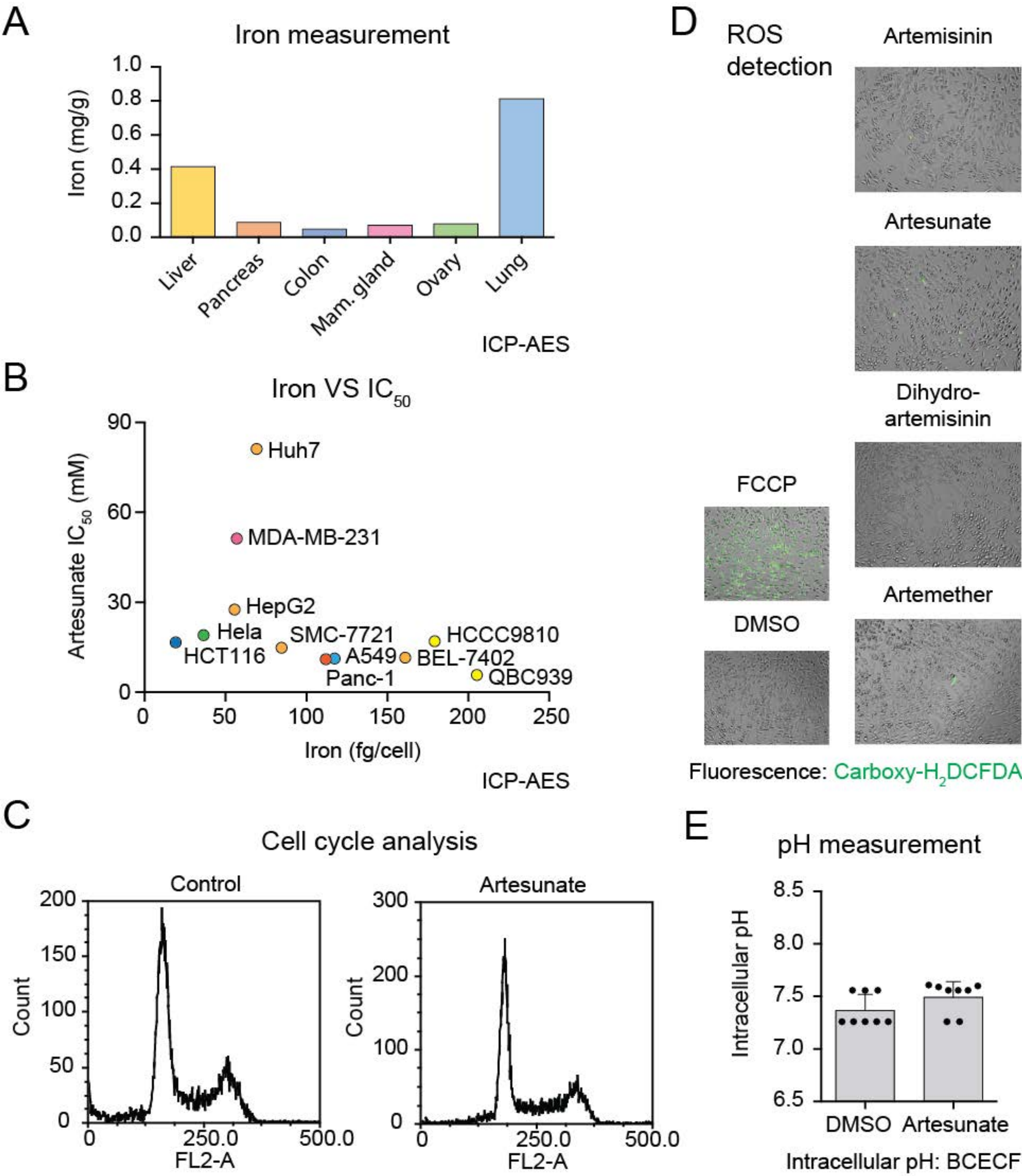

### Supplementary Figure 4

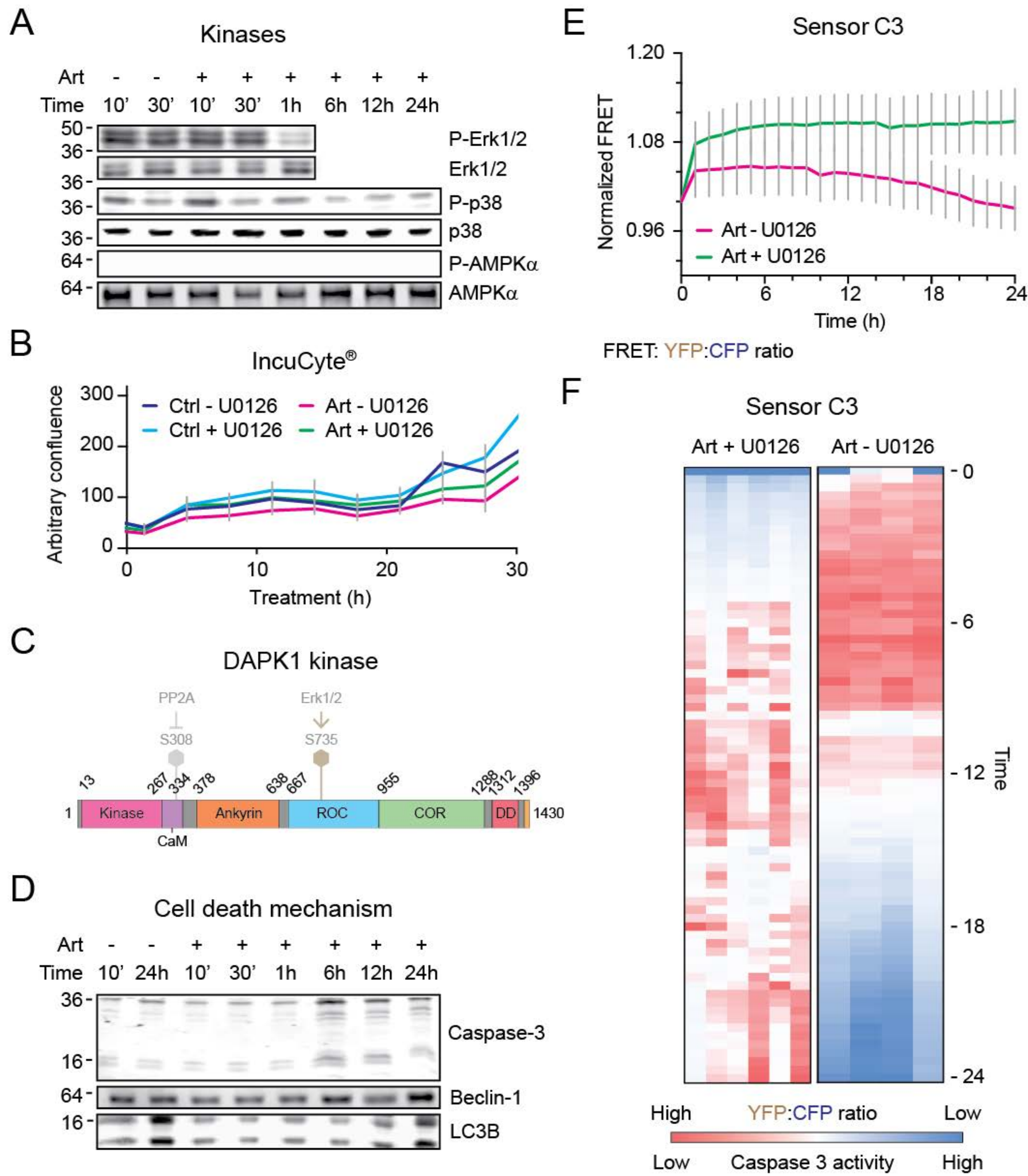

### Supplementary Figure 5

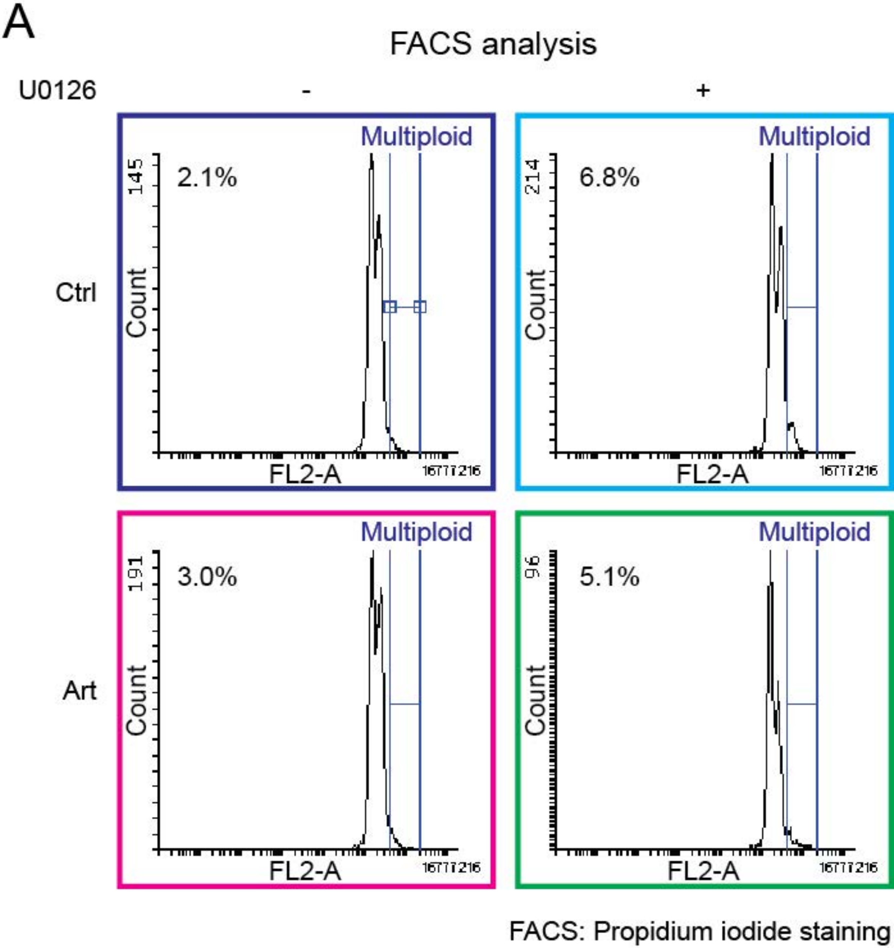
